# Muscle regeneration can be rescued in a telomerase deficient zebrafish model of ageing by MMP inhibition

**DOI:** 10.1101/2025.04.25.650659

**Authors:** Yue Yuan, Carlene Dyer, Robert Knight

## Abstract

Ageing progressively impairs skeletal muscle regeneration, contributing to reduced mobility and quality of life in the ageing population. Whilst the molecular changes underlying muscle ageing have been well characterised, their impact on muscle stem cell (muSC) behaviour during regeneration remains poorly understood. Here, we leverage the telomerase-deficient *tert* mutant zebrafish larvae as an *in vivo* model of accelerated ageing to perform real-time analysis of muSC dynamics following muscle injury. We demonstrate that the ageing-like inflammatory environment in *tert* mutant disrupts muSC migration, impairs activation and proliferation, and compromises regenerative capacity. We further show that sustained inflammation, mediated by persistent macrophage presence and elevated matrix metalloproteinase (MMP) activity directly limits muSC recruitment and migration efficiency. Pharmacological inhibition of MMP9/13 activity and genetic depletion of macrophages partially restore muSC migratory behaviour and regenerative outcomes. Notably, we demonstrate that muSC migration dynamics correlate with regenerative success, providing a functional readout for therapeutic screening. Our findings reveal zebrafish *tert* mutants offer a tractable system for dissecting age-associated changes to cell behaviour and for identifying rejuvenation interventions.

## Introduction

Skeletal muscle regeneration declines significantly with age, leading to impaired repair and progressive loss of muscle mass and function. This regenerative failure arises from both intrinsic muSC dysfunction and extrinsic changes in the tissue microenvironment (1, 2). In young muscle, injury triggers a well-orchestrated response in which muSC activate, proliferate, and differentiate into new myofibres, while immune cells facilitate repair through transient inflammatory signalling. However, ageing disrupts this balance. Aged muSCs exhibit reduced proliferative capacity and increased susceptibility to senescence, driven by chronic activation of stress pathways such as p38 MAPK signalling (3). Notably, while p38 MAPK activity is crucial for muSC differentiation under normal conditions (4, 5), its persistent activation in aged tissues contributes to impaired regeneration by prematurely limiting muSC proliferation and self-renewal (6). This cell-intrinsic deterioration is compounded by changes in the aged muscle niche, where chronic low-grade inflammation (“inflammaging”) leads to prolonged exposure to inflammatory cytokines such as IL-6 and TNFa. Overactivation of NF-κB signalling, a key transcriptional regulator of pro-inflammatory pathways, is also implicated in various age-related diseases, including sarcopenia (7–9). This inflammatory persistence drives muSC exhaustion and impairs differentiation, contributing to fibrotic tissue remodelling and inefficient muscle repair (10).

Although a number of studies have examined the impact of ageing on muSC function, we know little about how ageing affects muSC behaviour during regeneration (1, 11, 12). In particular, it remains unclear whether aged muSCs are less efficient at migrating to injury sites or have altered interactions with their environment and other cell types, such as macrophages, that are crucial for orchestrating repair. To investigate how ageing affects muSC function, we employed zebrafish (Danio rerio) as a vertebrate model and utilised telomerase-deficient (*tert* mutant) zebrafish larvae to study muscle regeneration in an ageing-like context. Zebrafish share key regenerative mechanisms with mammals but offer unique advantages for live imaging of cellular dynamics (13). The *tert* mutant zebrafish model provides a system of premature ageing driven by telomere dysfunction, leading to tissue atrophy, chronic inflammation, and impaired regenerative capacity (14–16). The recent study has confirmed that telomerase dysfunction during larval stages shows accelerated ageing, characterised by increased cellular senescence, elevated apoptosis and tissue degeneration (17). While *tert* mutants do not fully recapitulate all features of mammalian muscle ageing, such as systemic metabolic and endocrine changes, they serve as a valuable model for studying how telomere-dependent ageing the inflammatory environment during muscle repair. Importantly, telomere attrition in zebrafish induces a persistent inflammatory state that parallels inflammaging in aged mammals, making it a suitable system to examine how chronic inflammation affects muSC function *in vivo* (15).

In this study, we show that *tert* mutants reproduce key hallmarks of aged muscle repair, exhibiting delayed muscle fibres restoration, reduced muSC proliferation, and persistent macrophage infiltration following injury. Live imaging further reveals that chronic inflammation in tert impairs muSC activation and migration, directly linking age-associated inflammatory changes to disrupted stem cell behaviour. By capturing these cellular dynamics in real time, this approach fills a critical gap in understanding how ageing alters muSC function and interactions in real-time tissue repair context. Together, our findings highlight the zebrafish *tert* mutant as a powerful model for visualising stem cell–immune cell interactions in an ageing-context and provide new insights into how chronic inflammation drives regenerative failure with age.

## Results

### A zebrafish *tert* mutant model reveals molecular and functional changes in zebrafish larvae muscle

We aimed to understand whether muscle in telomerase deficient (*tert* mutant) zebrafish larvae show ageing like changes. To profile muscle, we performed RNA sequencing (RNA-seq) on dissected muscle from tert mutant and WT zebrafish larvae at 5 days post-fertilisation (5 dpf) (Figure 1A). Differential gene expression and Gene Ontology (GO) analyses revealed significant changes in genes related to inflammation, matrix organisation, cellular stress, and proteostasis (Figure 1B–C). Prominent upregulation of inflammatory markers such as *mmp9, cxcl18b*, and *tnfrsf1a* indicated an elevated inflammatory state in *tert* mutant muscle.

**Fig. 1.**
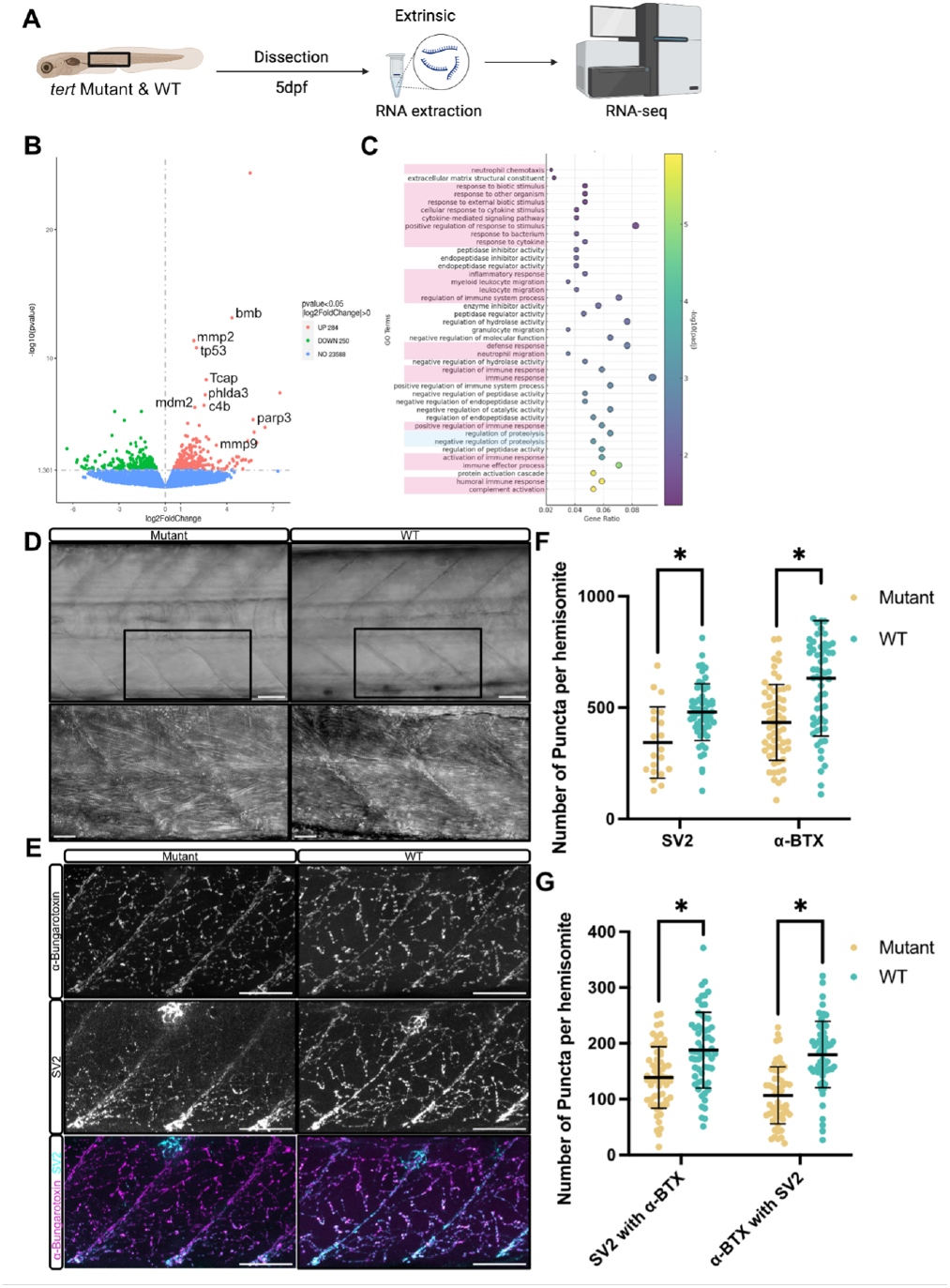
Muscle of zebrafish *tert* mutant larvae show anatomical and molecular changes reminiscent of ageing. A) Schematic representation of the experimental workflow. Muscle tissue was dissected from *tert* mutant and WT zebrafish larvae at 5 dpf, followed by RNA extraction and RNA-seq analysis. (B) Volcano plot of differentially expressed genes in *tert* mutant muscle compared to WT controls. Up-regulated genes involved in inflammation (*tp53, bmb*), DNA damage (*tp53, parp3*) and extracellular matrix (ECM) remodelling (*mmp2, mmp9*) are highlighted. (C) Gene ontology (GO) enrichment analysis of differentially expressed genes in *tert* mutant muscle. Terms related to inflammation and stress response highlighted in pink and loss of proteostasis highlighted in blue. (D) Brightfield images of muscle morphology in WT and *tert* mutant larvae. (E) Representative immunofluorescence images of NMJs labelled with α-bungarotoxin (AChR clusters, cyan) and synaptic vesicle protein SV2 (presynaptic marker, white). (F) Quantification of SV2 and α-bungarotoxin puncta in ventral myotomes of *tert* mutants and WT siblings. (G) Quantification of colocalised puncta between α-bungarotoxin and SV2. Number of animals used n = 8 (*tert* mutant), n = 7 (WT). Data are presented as mean ± SD and statistical significance was determined using unpaired Student’s t-test (* p < 0.05). Scale bars: 50 µm.

Histological examination of muscle tissue revealed no obvious morphological differences between *tert* mutant and WT larvae. Muscle fibres in both genotypes appeared structurally intact and well-aligned (Figure 1D). However, neuromuscular junction (NMJ) integrity, assessed through immunofluorescent staining of presynaptic marker SV2 and postsynaptic acetylcholine receptors (AChR, labelled with α-bungarotoxin), was significantly compromised in *tert* mutants. Quantitative analysis revealed a reduced number of individual presynaptic and postsynaptic puncta in *tert* mutant larvae (Figure 1E-F). Moreover, the number of colocalised puncta was significantly reduced in *tert* mutants, indicating impaired synaptic connectivity and possible denervation (Figure 1G). In addition, we observed reduced muSC proliferation using BrdU incorporation assays, suggesting impaired regenerative potential in *tert* mutant larvae (Supplementary Figure S1).

Together, these findings show that loss of *tert* function induces molecular signatures associated with inflammation and stress, accompanied by impaired neuromuscular connectivity, despite normal overall muscle fiber morphology. Common phenotypes between those we observe in muscle of *tert* mutant larvae and those described in ageing mammals suggest an accelerated ageing regime of muscle in zebrafish *tert* mutant muscle.

### Loss of tert function impairs muscle stem cell activation, proliferation and regeneration

We wished to know how the molecular changes observed in *tert* mutants related to muSC function during regeneration (Figure 2A). Following needle stick injury of the muscle, we quantified muSC activation at 24 hpi using Pax7 expression and found *tert* mutant larvae showed significantly fewer Pax7^+^ muSCs compared to WT (Figure 2B–C), indicating a defective activation of muSCs in response to injury.

**Fig. 2.**
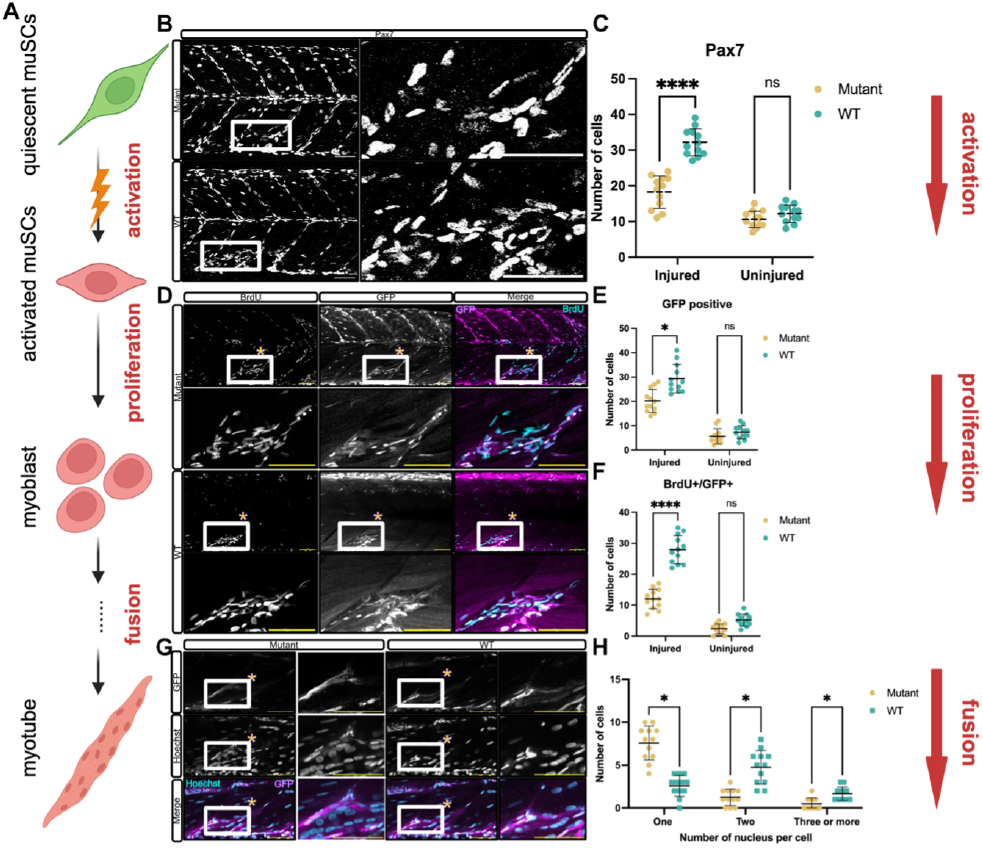
Loss of tert impairs muSC activation, proliferation and regeneration. (A) Schematic representation of muSC activation, proliferation, and fusion process following injury. Quiescent muSCs become activated, proliferate, and ultimately fuse to form multinucleated myofibres during regeneration. (B) Representative images showing Pax7 staining in WT and *tert* mutant larvae at 24 hpi. (C) Quantification of GFP positive cells in *tert* mutant and WT. Number of animals used n =12 (*tert* mutant), n=12 (WT). (D) Representative images showing BrdU incorporation (cyan) and GFP expression (magenta) in larvae expressing a pax7a:egfp transgene at 24 hpi. (E-F) Quantification of GFP and BrdU/GFP double-positive cells in *tert* mutant larvae and WT following injury. Number of animals used n =12 (*tert* mutant), n=12 (WT). (G) Representative images of regenerating muscle at 96 hpi, showing multinucleated myofibres in WT and *tert* mutant larvae. Hoechst staining (blue) marks nuclei. (H) Quantification of multinucleated myofibres. Number of animals used n =12 (*tert* mutant), n=12 (WT). Data are presented as mean ± SD and statistical significance was determined using an unpaired Student’s t-test (ns not significant, * p < 0.05, **** p < 0.0001). Scale bars: 50 µm.

To determine whether this corresponded to a change of muSC proliferation, we used larvae expressing a pax7a:egfp transgene to identify muSC-derived cells and measured BrdU incorporation after injury. We noted there were fewer GFP positive cells in *tert* mutants and these cells showed lower incorporation of BrdU by 24 hpi compared to WT (Figure 2D–F). Importantly, this difference was not attributed to increased cell death in *tert* mutants, acridine orange staining did not reveal discernible differences in cell death relative to WT larvae. (Supplementary Figure S2).

To understand whether regeneration of multinucleate myofibres was affected in *tert* mutants, we quantified nuclear number in newly formed GFP positive myofibres. Quantification of myofibre nuclear number revealed fewer multinucleated muscle fibres in *tert* mutant larvae at 96 hpi, indicating reduced myoblast fusion (Figure 2G–H). Collectively, these findings reveal that an impaired response by muSCs to injury occurs in tert mutants, resulting in attenuated regeneration of muscle.

### muSC behaviour during regeneration are disrupted in *tert* mutants

Given the critical role of muSC migration for muscle regeneration, we asked whether changes to the capacity of muSCs to respond to injury in tert mutants corresponded with an altered migratory behaviour. Using live imaging of larvae expressing the pax7a:egfp transgene, we tracked muSC movements from 6 hpi to 24 hpi in both *tert* mutant and WT larvae (Figure 3A). Quantification of muSC localisation and movement revealed striking differences between WT and *tert* mutant larvae (Figure 3B-G). In WT larvae, muSCs migrated efficiently toward injury sites and accumulated progressively, becoming aligned with the injury by 24 hpi. Conversely, muSCs in *tert* mutants exhibited a reduced and restricted migration, with fewer cells present at the injury over time (Figure 3A–C).

**Fig. 3.**
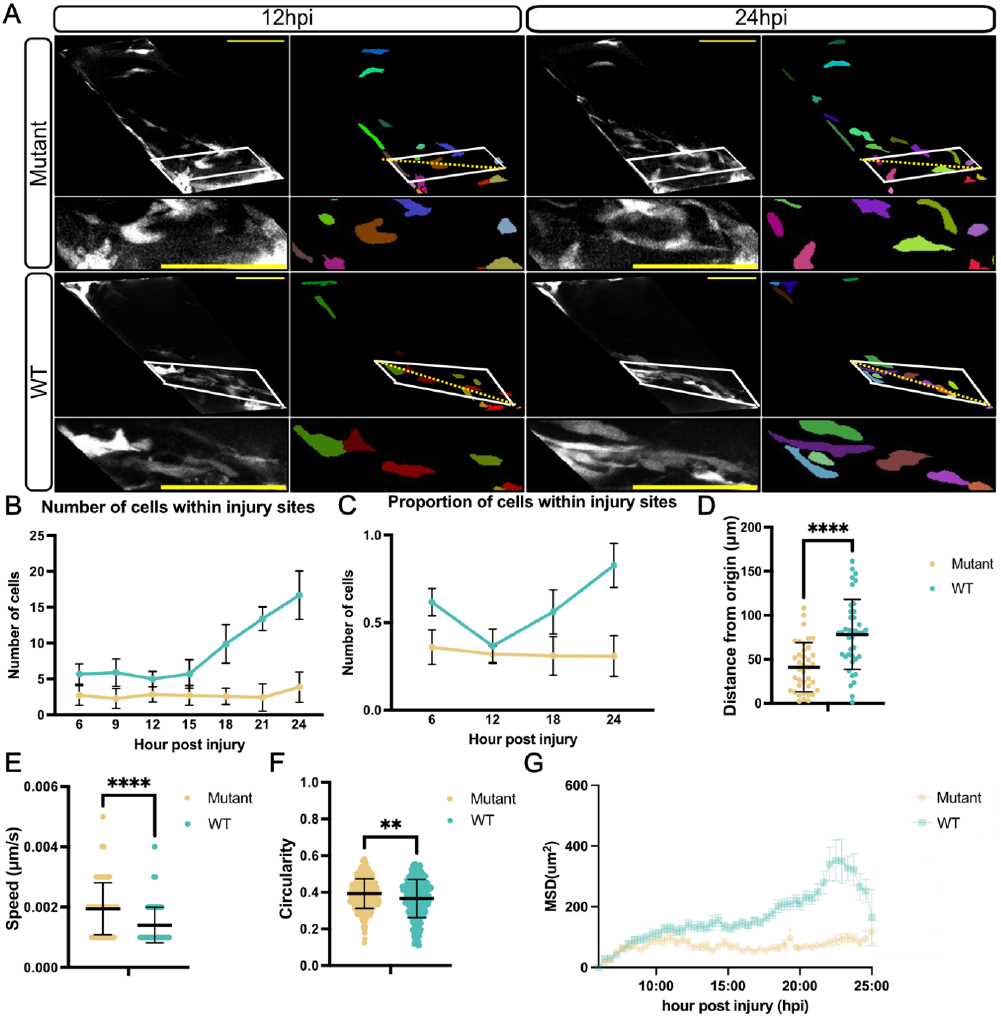
muSC migration is altered in *tert* mutants. (A) Images of time-lapsed movies of muscle from 6 hpi to 24 hpi in WT and *tert* mutant larvae expressing pax7a:egfp. Lower panels show segmented cells (coloured). Yellow dash line indicates the injury site. Quantitative analyses show (B) the number of muSCs within injury sites, (C) the proportion of cells within the injury sites relative to total muSC numbers in the injured myotome, (D) distances travelled from origin, (E) mean cell speed, (F) cell circularity, and (G) mean squared displacement (MSD). Number of animals used n = 7 (*tert* mutant), n = 10 (WT). Data are presented as mean ± SD and statistical significance was determined using an unpaired Student’s t-test (** p < 0.01, **** p < 0.0001). Scale bars: 50 µm.

Further detailed analysis of cell movement and cell shape revealed that although mean cell speed was increased, muSCs moved significantly shorter distances from their origin and showed greater circularity, indicative of fast yet confined movements (Figure 3D–F). Multiple linear regression models were generated to test whether the mean squared displacement (MSD) showed significant differences between tert mutant and WT. MSD was significantly lower in *tert* mutants than in WT animals, indicative of impaired cell migration (Figure 3G).

Our characterisation of cell response to injury reveals an abnormal migratory response by muSCs in *tert* mutants, characterised by an increased speed but showing reduced directional movement, resulting in fewer cells present within the injury at 24 hpi.

### Injured *tert* mutants exhibit prolonged inflammation and delayed macrophage clearance following injury

To understand whether *tert* mutants show an altered response to injury that affects muSC responses, we performed RNA-seq on injured muscle tissue of 5 dpf larvae at 24 hours after needle-stick injury (Figure 4A). KEGG pathway analysis revealed significant upregulation of inflammatory pathways in tert mutants compared to WT larvae, particularly those associated with ECM remodelling and immune response (Figure 4B). Among these, we identified a strong upregulation of MMP genes in injured *tert* mutant muscle compared to injured WT (Figure 4C). Using qPCR and HCR we confirmed elevated expression of *mmp9* and *mmp13* in *tert* mutant muscle following injury (Figure 4D,E).

**Fig. 4.**
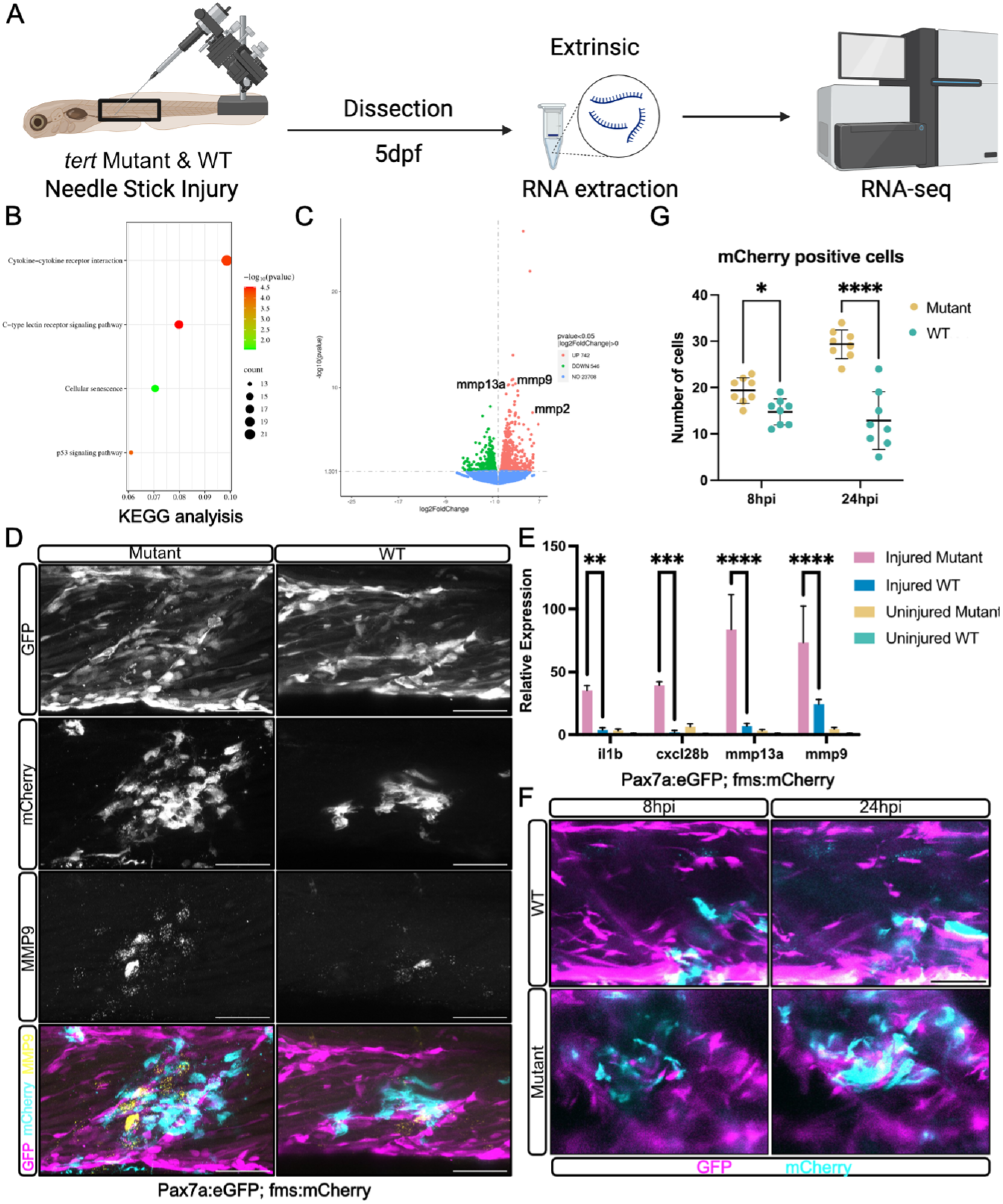
Injured *tert* mutant shows prolonged inflammatory response and delayed macrophage clearance following injury. (A) Schematic representation of the experimental workflow. Muscle from injured *tert* mutant and WT zebrafish larvae was extracted and profiled by RNA-seq. (B) KEGG pathway analysis of differentially expressed genes in *tert* mutant vs WT injured muscle. (C) Volcano plot showing upregulation of *mmp2/mmp9/mmp13* in *tert* mutant injured muscle. (D) Representative images of HCR detection of mmp9 expression relative to muSCs expressing pax7a:egfp (GFP, magenta) and macrophages expressing fms:mCherry (mCherry, cyan); number of animals n = 6 (*tert* mutant), n = 6 (WT). (E) qPCR measures of inflammatory (*il1b, cxcl28b*) and *mmp13a, mmp9* gene expression. (F) Representative images of muSCs expressing pax7a:egfp (GFP, magenta) and macrophages expressing fms:mCherry (mCherry, cyan) in injured muscle of *tert* mutant and WT larvae at 8 hpi and 24 hpi. (G) Quantification of macrophages in *tert* mutants and WT larvae at 8 hpi and 24 hpi, number of animals used n = 10 (*tert* mutant), n = 10 (WT). Data are presented as mean ± SD and statistical significance was determined using an unpaired Student’s t-test (* p < 0.05, ** p < 0.01, *** p < 0.001, **** p < 0.0001). Scale bars: 50 µm.

Given that the regenerating muscle of *tert* mutants shows a strong inflammatory signature, which may perturb immune cell function, we examined macrophage responses to injury. In WT larvae, macrophages were efficiently cleared from the injury site by 24 hpi, while in *tert* mutants, macrophages persisted at the site of injury, indicating delayed clearance (Figure 4F). Quantification of mCherry positive macrophages revealed significantly higher numbers in *tert* mutants at both 8 hpi and 24 hpi, further supporting the notion of prolonged inflammation in tert mutants (Figure 4G).

### MMP9/13 inhibitor I treatment restores muSC activation, proliferation and migration in *tert* mutants

Metal-loproteinase are major regulators of tissue remodelling and are expressed by macrophages during muscle injury (18). Both *mmp9* and *mmp13* were significantly upregulated in injured *tert* mutant muscle compared to WT muscle suggesting they may be responsible for the impaired muSC response. We therefore tested how MMP inhibition affects muSC responses to injury by treating *tert* mutant and WT larvae with MMP9/13 inhibitor I 24 hours prior to injury and subsequently after injury then examined muSC responses. Relative to untreated *tert* mutants, the number of Pax7 positive muSCs was significantly restored in *tert* mutants, reaching levels comparable to WT controls (Figure 5A, B). Similarly, proliferation of muSCs expressing pax7a:egfp was also significantly improved in inhibitor treated *tert* mutants compared to untreated *tert* mutants (Figure 5C, D, E). These results suggest that excessive MMP9/13 activity impairs muSC activation and proliferation, and its inhibition rescues these early regenerative processes.

**Fig. 5.**
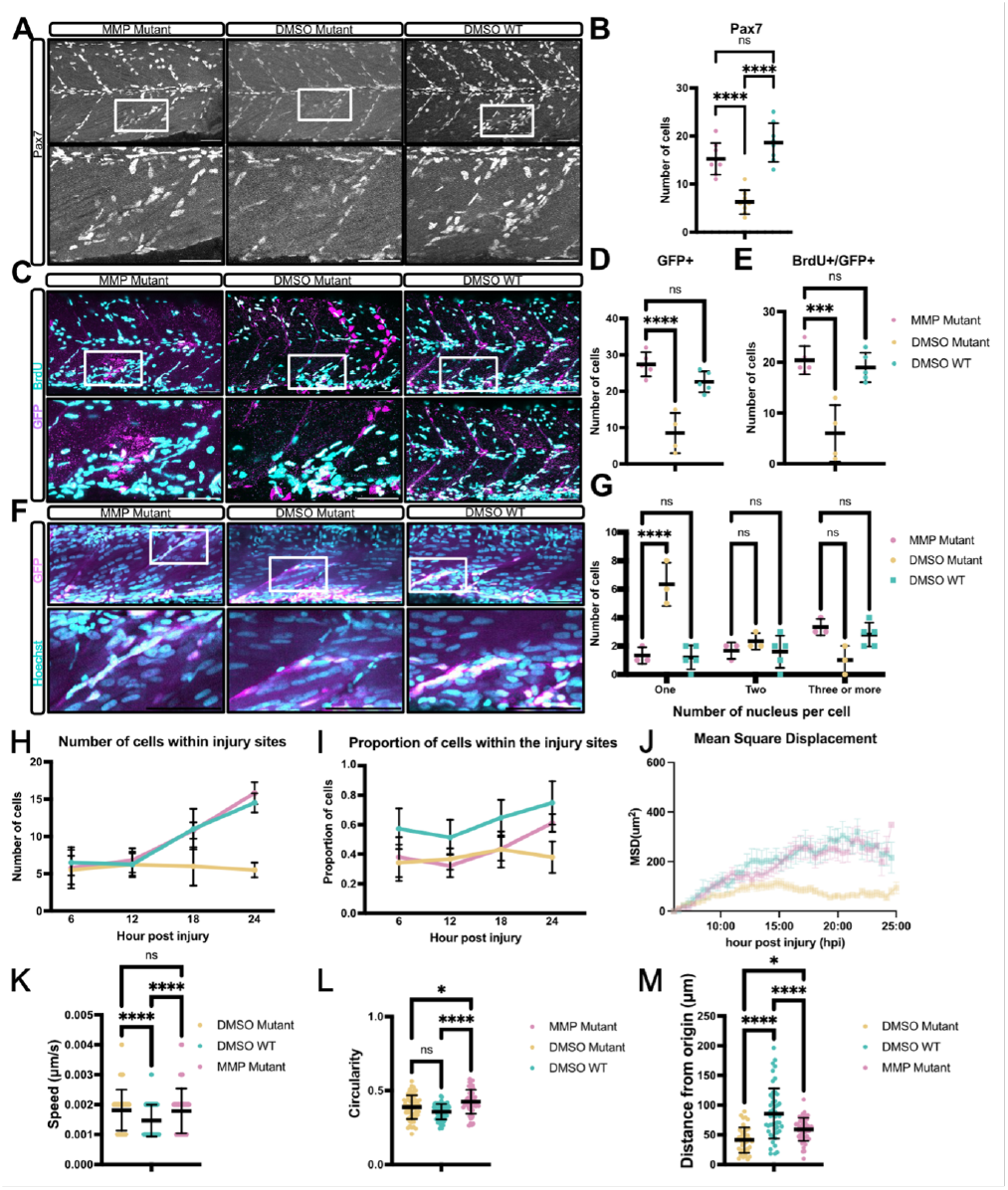
MMP9/13 inhibitor I treatment restores muSC activation, proliferation and migration in *tert* mutants. (A) Immunolabelling of Pax7 in *tert* mutant and WT larvae treated with MMP9/13 inhibitor I. Number of animals used n = 10 (*tert* mutant + inhibitor), n = 8 (*tert* mutant + DMSO), n = 6 (WT + DMSO). (B) Quantification of muSCs expressing pax7a:egfp in muscle of injured *tert* mutant treated with MMP9/13 inhibitor compared to untreated *tert* mutant and WT. (C) Representative images of BrdU labelling (cyan) relative to muSCs expressing pax7a:eGFP (GFP, magenta). Number of animals used n = 12 (*tert* mutant + inhibitor), n = 9 (*tert* mutant + DMSO), n= 7 (WT + DMSO). (D-E) Quantification of GFP positive (magenta) muSCs and muSCs expressing both BrdU (cyan) and GFP (magenta) in inhibitor treated *tert* mutant larvae and control groups. (F) Representative images of multinucleated myofibres following inhibitor treatment. (G) Quantification of multinucleated myofibres. Number of animals used n = 9 (*tert* mutant + inhibitor), n = 6 (*tert* mutant + DMSO), n= 6 (WT + DMSO). Quantification of the number (H) and the proportion (I) of muSCs accumulating at the injury site over time in inhibitor treated *tert* mutant and control groups. (J) MSD analysis of muSC movement in treated *tert* mutant compared to untreated *tert* mutant and WT animals. (K-M) Quantification of muSC migration parameters, including mean cell speed (K), mean cell circularity distance travelled from origin (M). Number of animals used n = 10 (*tert* mutant + inhibitor), n = 8 (*tert* mutant + DMSO), n= 8 (WT + DMSO). Data are presented as mean ± SD and statistical significance was determined using an unpaired Student’s t-test (ns not significant, * p < 0.05; *** p < 0.001, **** p < 0.0001). Scale bars: 50 µm.

To evaluate whether this improvement in stem cell function translates into improved muscle regeneration, we quantified multinucleated myofibres at 96 hpi. Inhibitor treated *tert* mutant larvae exhibited a significant increase in the number of multinucleated fibres, indicating enhanced fusion and improved regeneration (Figure 5F, G). Further examination of myofibre morphology at 6 dpi using phalloidin revealed alignment of the newly regenerated fibres was similar to that seen in WT larvae (Supplementary Figure S3A).

These results demonstrate that excessive MMP9/13 activity contributes to the impaired regenerative phenotype in *tert* mutants, and inhibition of MMPs can effectively rescue muSC function and muscle regeneration.

### Partial restoration of muSC migration by MMP9/13 inhibition

In *tert* mutants, muSCs showed a reduced proliferative response to injury, which coincided with an altered migratory behaviour. To understand whether rescue of muSC proliferation, differentiation and fusion by MMP9/13 inhibition corelated with a rescue of cell behaviour we tracked muSC migration from 6 hpi to 24 hpi in larvae treated with MMP9/13 inhibitor I and compared to DMSO treated controls (Supplementary Figure S4).

Quantitative analyses showed that MMP9/13 inhibition significantly increased the absolute number of muSCs reaching the injury site in *tert* mutants compared to untreated *tert* mutants (Figure 5H). However, the proportion of muSCs within the injury site, relative to total muSC numbers, is consistently smaller than that in the DMSO treated controls (Figure 5I). Furthermore, analysis of migration trajectories revealed that inhibitor treated *tert* mutant muSCs displayed significantly greater displacement from their origin compared to untreated *tert* mutant cells, indicative of partially restored migratory capacity (Figure 5J).

Despite observing a similar number of cells around the injury site (Figure 5H, I), as well as comparable MSD (Figure 5J) and mean cell speed (Figure 5K) between treated *tert* mutants and WT controls, detailed analysis revealed that muSCs from inhibitor treated *tert* mutants still exhibited different migratory behaviours relative to WT animals. Specifically, there was proportionately fewer cells reaching the injury site (Figure 5I), they showed a shorter migratory distance from their origin (Figure 5M) and showed increased cell circularity compared to WT muSCs (Figure 5L). These observations indicate that although overall migratory behaviour and accumulation at the injury site were improved, certain aspects of the migratory pattern remained abnormal.

### Macrophage depletion restores muscle stem cell activation and proliferation in *tert* mutants

To understand whether macrophage behaviour was affected by MMP9/13 inhibitor I treatment, we performed live imaging and compared against untreated control *tert* mutants. We observed a general decrease of the number of macrophages in the injured myotome of inhibitor treated *tert* mutants compared to the DMSO treated control (Supplementary Figure 5). To better understand whether macrophage presence directly impacts muSC behaviour and proliferation in an ageing-like context, we utilised the Metronidazole-Nitroreductase (MTZ-NTR) system to perform macrophage depletion (19). Successful macrophage depletion was confirmed via fluorescence microscopy (Supplementary Figure S6).

Following macrophage depletion, we observed a significant restoration in the number of Pax7 positive muSCs and an increase in BrdU incorporation by muSCs, indicating improved activation and proliferation of muSCs in *tert* mutants (Figure 6A, B, D-F). However, analysis at 6 dpi revealed persistent abnormalities in myofibre organisation in *tert* mutants following macrophage depletion (Supplementary Figure S3B). To determine whether re-innervation had been rescued, NMJ formation in regenerating muscle was quantified. An increased number of presynaptic and postsynaptic puncta were detected at the injury site in *tert* mutants lacking macrophages, but this was not comparable to WT larvae (Figure 6C, G–H). This suggests that removing macrophages in *tert* mutants did not completely rescue regeneration of muscle.

**Fig. 6.**
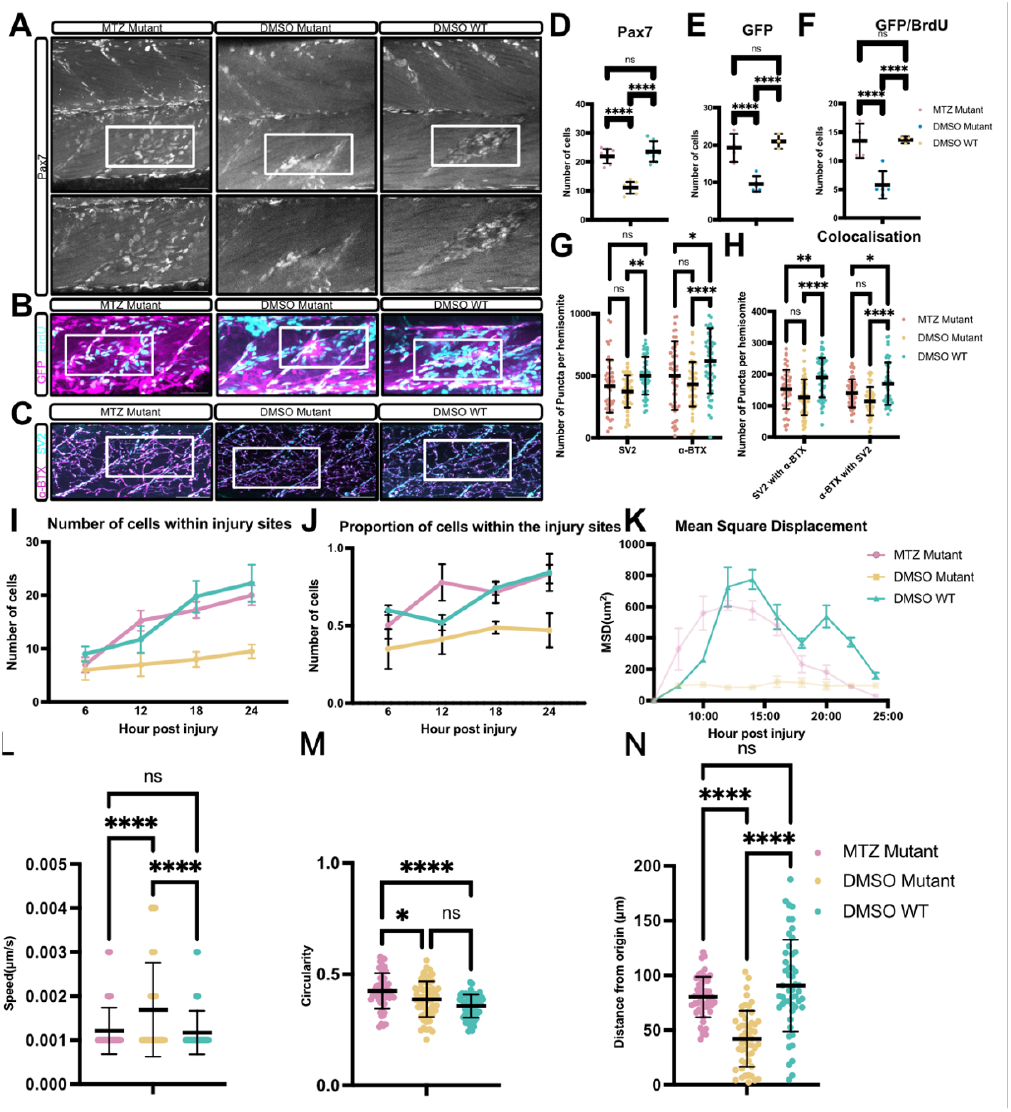
Macrophage depletion restores early muSC responses but does not rescue late stage muscle regeneration in *tert* mutant larvae. (A) Representative images of Pax7 staining in *tert* mutant and WT larvae with and without MTZ treatment. Number of animals used n = 8 (all conditions). (B) Representative images of BrdU (cyan) incorporation in larvae expressing pax7a:egfp (GFP, magenta). Number of animals used n = 4 (*tert* mutant + MTZ), n = 5 (*tert* mutant + DMSO), n = 3 (WT + DMSO). (C) Representative images of NMJs at 6 dpi using α-bungarotoxin (α-BTX; magenta) to detect AChR and SV2 (cyan) immunostaining to detect pre-synaptic vesicles. Number of animals used n = 8 (all conditions). (D) Quantification of GFP positive muSCs in MTZ treated *tert* mutant larvae expressing pax7a:egfp and DMSO treated *tert* mutants. (E, F) Quantification of BrdU and BrdU/GFP double-positive cells in MTZ treated *tert* mutant larvae expressing pax7a:egfp and DMSO treated *tert* mutants. (G, H) Quantification of SV2 or α-BTX puncta, and puncta showing colocalisation in muscle in larvae following macrophage depletion. Number of animals used n = 8 (*tert* mutant + MTZ), n = 6 (*tert* mutant + DMSO), n = 7 (WT + DMSO). Quantitative analyses show (I) the number of muSCs expressing pax7a:egfp at the injured muscle, (J) the proportion of cells within the injury sites relative to total muSC numbers within the myotome, (K) mean squared displacement (MSD), (L) mean cell speed, (M) cell circularity and (N) distances travelled from origin by muSCs. Number of animals used n = 6 (*tert* mutant + MTZ), n = 4 (*tert* mutant + DMSO), n = 5 (WT + DMSO). Data are presented as mean ± SD and statistical significance was determined using an unpaired Student’s t-test (ns not significant, * p < 0.05, ** p < 0.01, **** p < 0.0001). Scale bars: 50 µm.

Together, these findings suggest that macrophage depletion partially restores early stem cell responses, but is insufficient to fully rescue structural and functional regeneration at later stages.

### Macrophage depletion alters muSC migration dynamics

Given that *mmp9* expression by macrophages affects muSC proliferation in a non-ageing context (18), we wished to understand whether the elevated *mmp9* expression and altered macrophage responses could explain the impaired muSC response to injury in *tert* mutants. To determine how macrophage depletion affected muSC migration dynamics during regeneration we visualised muSCs expressing pax7a:egfp (Supplementary Figure S7. We found that macrophage-ablated *tert* mutants exhibited earlier and more robust accumulation of muSCs at the injury site compared to untreated *tert* mutants, with a peak in cell number occurring earlier in the regeneration process (Figure 6I). This trend was also reflected in the proportion of muSCs localised within the injury zone, which increased more rapidly and declined earlier in the MTZ treated *tert* mutants, suggesting an earlier recruitment of muSCs in an absence of macrophages (Figure 6J).

Further analysis of migration characteristics showed that macrophage depletion partially restored several aspects of migratory behaviour. Treated *tert* mutants showed a significant increase in distance travelled from the point of origin (Figure 6N) and a reduction in cell circularity (Figure 6M), reflecting more efficient and directional movement. However, mean cell speed remained elevated in MTZ treated *tert* mutants (Figure 6L), and most notably, mean squared displacement (MSD) remained significantly different than in controls (Figure 6K).

These results indicate that macrophage depletion enhances initial muSC recruitment and partially rescues migratory behaviour in an ageing-like context. However, the persistent deficit in MSD highlights those additional mechanisms, beyond macrophage presence alone, contribute to impaired cell dynamics in *tert* mutant muscle.

## Discussion

Our study reveals that an ageing-like inflammatory environment is refractory to muscle regeneration in *tert* mutants, accompanied by impaired muSC activation and migratory dynamics. This is mediated by an inappropriate activity of metalloproteases secreted by macrophages, which inhibit muSC migration and generation of new myofibres. By deploying *in vivo* imaging, we demonstrate that the dynamics of muSC responses to injury can be linked to their functional capacity to repair muscle, linking cellular behaviour to regenerative outcomes. These findings underscore how chronic inflammatory and ECM changes to the muscle environment affect muSC function *in vivo*, providing new insight into the basis for impaired muscle regeneration in ageing.

Telomerase-deficient zebrafish rapidly develop an ageing-like phenotype, making them a powerful model to study how age-related changes in the tissue environment impact regeneration. Telomere shortening in *tert* mutant triggers increased cellular senescence and chronic, systemic inflammation (17, 20, 21). In this study, we observed molecular changes akin to aged muscle in zebrafish *tert* mutant larvae, involving upregulation of stress responsive gene expression and an exaggerated inflammatory response. Similar to ageing in mice, muSCs in *tert* mutants have a delayed activation and proliferation response after injury, and often a fraction of muSC pool becomes senescence (22). The *tert* mutant larvae recapitulate this age-associated decline in function, evidenced by reduced muSC activation and proliferation after muscle damage. We also observe elevated expression of senescence markers *p16* and *p21* in *tert* mutant muscle, indicative of a senescent program, reminiscent of aging mammalian muscle.

Crucially, our live imaging experiments have demonstrated how ageing may affect the interaction between muSCs and immune cells in a regeneration context. After injury or damage, macrophages infiltrate to phagocytose tissue, activate resident stem cells and then mostly exit the injury site allowing muSCs to proceed with regeneration. In *tert* mutants, we observed macrophages lingering at injury sites well beyond the acute phase and physically clustering around muSCs. This persistent macrophage presence correlated with muSCs retaining a rounded, non-spreading state, failing to adopt the spreading morphology typically associated with effective migration and differentiation. Such rounded cell shapes have previously been described for migratory cells experiencing altered ECM conditions or physical constraints, as reported in studies of macrophages and neutrophils during ECM remodelling (23, 24). Similarly, ECM composition and stiffness have been shown to significantly influence cell shape and migration patterns, as changes in ECM stiffness guide cells toward injury sites and modulate their morphology and speed of movement (24, 25). Furthermore, inflammatory cells may actively restraining muSCs, either through direct cellcell contact or continual exposure to inflammatory cytokines (23). Recent intravital imaging studies in mice have begun to characterise muSC interactions with macrophages under disease condition through intravital imaging, demonstrating prolonged contacts between muSCs and macrophages, which negatively impact on stem cell motility and regenerative outcomes (26). In the study, muSC and macrophage interactions during muscular dystrophy are often characterised by macrophage derived factors restricting muSC migration and differentiation, leading to impaired regeneration. Our zebrafish analyses extend these observations by muSCs in an ageing-like context, reinforcing the concept that an effective regenerative response requires an acute, transient inflammatory reaction, followed by timely resolution to allow the initiation of regeneration.

In *tert* mutants, we observed elevated expression of MMP2/9/13 genes after injury, similar to descriptions of elevated MMP9 expression in muscle of aged mice (27). MMPs are well-established regulators of muscle injury repair, orchestrating both immune cell infiltration and tissue remodelling. In young mice, inflammatory cells (such as neutrophils and macrophages) secrete MMPs like MMP9 early after muscle injury to degrade components of the basement membrane, allowing leukocytes to rapidly infiltrate damaged tissue (28, 29). This proteolytic clearance not only removes debris but also creates space for incoming cells and releases ECM bound signals that further recruit inflammatory cells. As regeneration progresses, other MMPs (e.g. MMP2 and MMP13) become active to remodel the ECM, which facilitates migrating muSCs to reach the injury and form new myofibres (18, 27, 28). Thus, in a young environment MMPs play a pro-regenerative role by balancing timely inflammation and creating a permissive scaffold for repair. Consistent with these roles, muscle injuries in MMP deficient models often show aberrant tissue repair. Inhibition or genetic ablation of MMP9 can delay macrophage infiltration and modify regeneration outcomes (29), whereas excessive or prolonged MMP activity can be deleterious, as seen by chronic high MMP9 activity in dystrophic muscle exacerbating tissue damage and inflammation (30). These observations suggest that precise regulation of MMP expression and activity is crucial for effective muscle regeneration.

Our findings point to two major extrinsic regulators of muSC behaviour in an ageing context: macrophage mediated inflammation and ECM remodelling. Both factors are known to influence muSC function in normal regeneration, and our analyses in telomerase deficient animals show how their dysregulation leads to defective outcomes. To determine how macrophage function contributes to this phenotype, we showed that ablation of macrophages in *tert* mutant partially restored muSC function and behaviour. However, we observed this resulted in altered cell dynamics with many more muSCs rapidly accumulating at the wound site earlier than observed in the untreated *tert* mutant animals. This is presumably because inhibitory signals or physical blockage from macrophages are no longer present to prevent muSCs migrating into the damaged muscle area. Despite arriving at the injury site earlier in an absence of macrophages, the muSCs then pause, maintaining a rounded shape, prior to differentiation and fusion. This pattern of “early gather, late integration” suggests that persistent macrophages in ageing interfere with the initial phase of muSC recruitment to injury.

Conversely, when MMP activity is inhibited in *tert* mutants, muSCs show a different response to injury which is more similar to WT animals, showing a more directional and faster migration. This behaviour also resembles the coordinated sequence of responses we have previously described for muSCs in larval muscle involving migration/ arrive at injury/ elongate (31). This suggests that MMP9/13 inhibition has restored sufficient cues from the surrounding environment to enable a more normal muSC movement. However, even though muSC arrival timing improved, the efficiency of regeneration remained low, evidenced by a smaller proportion of muSCs at the injury site in *tert* mutants compared to WT. Differences in the type of behavioural changes to muSCs in *tert* mutants in response to macrophage removal or MMP inhibition suggests these perturbations are affecting different cell responses. Macrophage depletion increased the speed of muSC recruitment to the injury, whereas MMP inhibition rescued normal migratory behaviour.

Based on these observations we propose that macrophages and ECM components play complementary but distinct roles in regulating muSC behaviour in ageing. Persistent macrophages continuously emit inflammatory signals and may physically hinder muSCs, thereby delaying their initial activation and migration to injuries. On the other hand, an altered ECM modelling creates a suboptimal migratory landscape for muSCs, affecting how efficiently they can travel. Effective muscle regeneration requires both timely inflammation resolution and proper ECM remodelling; ageing skews both processes, and each must be addressed to rejuvenate regenerative capacity.

These findings advance our understanding of muscle ageing by demonstrating that a combination of niche-extrinsic ageing changes leads to regenerative failure. Using *in vivo* imaging, we reveal how muSC function is driven by a dynamic response to macrophage-dependent cues, which are impaired in ageing due to elevated MMP activity. We also highlight that the zebrafish *tert* mutant recapitulates many of the changes observed in ageing muscle and offers a powerful system for identifying ageing-associated factors driving the aberrant behaviour of distinct cell populations in tissue homeostasis and repair.

## Acknowledgements

We thank the aquarium facility at Kings College London for care and maintenance of fish lines and technical team in CCRB for support with imaging and analyses. We thank Dr. Miguel Godinho Ferreira for insightful discussions and advice regarding use of telomerase mutants and Dr. Elisabeth Busch-Nentwich for the telomerase mutant line.

## Author contribution

**Investigation**, Carlene Dyer, Yue Yuan; **Data curation**, Yue Yuan; **Formal analysis**, Carlene Dyer, Yue Yuan; **Funding acquisition**, Robert Knight; **Methodology**, Yue Yuan, Robert Knight; **Conceptualisation**, Yue Yuan, Robert Knight; **Visualisation**, Yue Yuan; **Supervision**, Robert Knight; **Writing – original draft**, Yue Yuan; **Writing – review & editing**, Robert Knight. All authors have read and agreed to the published version of the manuscript.

## Online Methods

### Fish Stocks and Husbandry

All animals used were reared at the Kings College London Zebrafish Facility and maintained in accordance with UK Home Office regulation, UK Animals (Scientific Procedures) Act 1986, under project licence PP9727122. Fish carrying allele *tert*^*sa6541*^ were obtained from Dr. Elisabeth Busch-Nentwich (1) and maintained in the AB background. Animals were in-crossed to generate *tert*^*sa654/1sa6541*^ mutants (*tert* mutants) or wildtype animals (WT) for experiments. Trangenic lines TgBAC[pax7a:egfp] (pax7a:egfp) (2) and TgBAC[fms:gal4; UAS:NfsB-mCherry] (fms:mCherry) (3) were crossed against animals carrying the *tert*^*sa65421*^ allele for visualisation of muSCs and macrophages. Fish were carefully monitored to prevent any suffering due to telomere attrition related phenotypes and only morphologically normal larvae with inflated swim bladders at 5 dpf entered the nursery to be raised to adulthood for this study. Embryos were obtained from natural spawning and embryonic fish were maintained in E3 Phenylthiourea (PTU) solution at 28.5°C to inhibit pigment formation medium (4).

### Genotyping

DNA from embryos or fin biopsies was extracted and genotyped for *tert* mutants using KASP genotyping as previous described (5).

### Needlestick injury

Larvae were anesthetised in 0.004% w/v Tricaine (Sigma-200mg/ml) in E3 media and mounted in 1.5% w/v low melt agarose (Sigma). To induce muscle injury, a sharpened tungsten wire was applied to the myotome as previously described (6). Injured larvae were then carefully released from the agarose and placed in multi-well dishes containing fresh E3 PTU.

### Immunolabelling and BrdU incorporation

The detection of Pax7 was performed as previous described (7). Briefly, 5 dpf larvae were euthanised with 0.4% w/v Tricaine and fixed in 2% w/v paraformaldehyde (PFA) for 45min and washed with 1% PBT (1XPBS, 1% v/v Triton X-100) before blocking in 5% goat serum for 1 hour at room temperature and incubated with Pax7 antibody mouse anti-Pax7 (developed by A. Kawakami at the Tokyo Institute of Technology, obtained from the Developmental Studies Hybridoma Bank (DSHB), created by the NICHD of the NIH and maintained at The University of Iowa, Department of Biology, Iowa City, IA, United States) for two nights. Samples were then washed in 1% PBT and incubated with Alexa conjugated secondary antibodies (Invitrogen) at 4°C overnight. Finally, samples were washed and post-fixed in 4% w/v PFA for 30 min at room temperature prior to addition of DAPI or Hoechst-33342 and mounting.

For other antibodies, larvae were euthanised with 0.4% w/v Tricaine and fixed in 4% w/v PFA overnight at 4°C. Larvae were washed in 0.1% PBT (1xPBS, 0.1% v/v Tween-20), followed by a methanol series (25, 50 and 75% v/v methanol in distilled H_2_O for 5min each), moved to 100% methanol at −20°C overnight. After a reverse methanol series, larvae were permeabilised in 10 µg/mL proteinase K (5dpf for 60min), rinsed and fixed at RT for 30min. Samples were then blocked in 10% v/v newborn calf serum (NBCS) for 1h at room temperature and incubated with primary antibody overnight at 4°C. Primary antibodies used included chick anti-GFP (ab13970, Abcam) and SV2 supernatant (Q7L0J3, DSHB). Acetylcholine receptors were detected using alpha Bungarotoxin CF555 (00018-100µg, Insight Biotechnology). Samples were then washed in 0.1% PBT and incubated with corresponding secondary antibodies (Invitrogen) at 4°C overnight before fixing in 4% w/v PFA for 30 min at RT prior to addition of DAPI or Hoechst-33342 and mount for imaging.

For the detection of BrdU, larvae were exposed to 10mM BrdU in the medium, following which they were fixed with 4% PFA and treated with 2M HCL for 1h at RT. HCL was subsequently neutralised by washing in 0.1M borate buffer (0.62g Boric acid, 1.5ml 75mM NaCl in 100ml water; pH8.5), then washed with PBT containing 1% v/v Triton X-100 and 1% v/v DMSO. Primary antibody used was rat anti-BrdU (ab6326, Abcam).

### Whole-mount immune-coupled hybridisation chain reaction (WICHCR)

WICHCR were carried out as described previously (8). Probes for *mmp9* were designed and purchased from Molecular Instruments.

### RNA isolation and qRT-PCR

Total RNA was isolated from dissected injured trunk regions of larvae at 24h post injury (hpi). Tissue was homogenised in TriReagent (Sigma) and total RNA isolated according to the manufacturer’s protocol. RNA concentration was measured using a Nanodrop spectrophotometer (Peqlab Biotechnology GmbH, ThermoFisher Scientific) and 2 µg, of total RNA was reverse transcribed into cDNA using random hexamer primers (Promega) and M-MLV reverse transcriptase (Promega).

The C1000 TM Thermal Cycler with CFX384 Optical Reaction Module (Bio-Rad) was used to perform quantitative real-time PCR experiments. Amplification after each PCR cycle was detected using TaqMan ™ Gene Expression Assays (Thermo Fisher Scientific) with biological and technical triplicates run on the same 384-well plates with a total volume of 5µl. Expression data were calculated according to the ΔΔCt method (9). Expression analysis is shown relative to the untreated controls and relative to 18s as internal expression control.

### RNA sequencing and analysis

All RNA analyses were performed using triplicates for all conditions using material isolated from the muscle of 30 larvae using Tri-reagent. mRNA was purified from total RNA using poly-T oligo-attached magnetic beads. After fragmentation, the first strand cDNA was synthesized using random hexamer primers, followed by the second strand cDNA synthesis using dTTP for non-directional library. Index of the reference genome was built using Hisat2 v2.0.5 and paired-end clean reads were aligned to the reference genome Danio_rerio.GRCz11.dna using Hisat2 v2.0.5 (10). Differential expression analysis was performed using the DESeq2R package (1.20.0) (11). Gene Ontology (GO) (12) enrichment analysis of differentially expressed genes was implemented by the ClusterProfiler R package, in which gene length bias was corrected.

### Treatment of larvae with chemicals

MMP9/13 inhibitor I (15942, Cayman Chemical) was reconstituted to a stock concentration of 20mM in fresh DMSO, aliquoted and stored at −80°C. The stock solution was then diluted to a working concentration in medium at a concentration of 100 µM.

Macrophage ablation was achieved by treating Tg-BAC[fms:gal4; UAS:NfsB-mCherry] (fms:mCherry) larvae with 5mM metronidazole (MTZ, Sigma) (13) in medium 24 hours prior to injury. Following injury, larvae were placed into fresh 5mM MTZ for the duration of the experiment. MTZ solution was refreshed every 12-14 hours.

### Imaging acquisition

Images of fixed samples were acquired using a Carl Zeiss LSM980 using a 20x air objective (NA 0.8). Z-stacks encompassing the total myotome (with 1µm Z-slices) were captured. Z-stacks were captured at a resolution of 1024 × 1024 pixels and each channel was averaged 4 times. For live imaging, larvae were anesthetised in 0.004% w/v Tricaine and mounted in 1.5% w/v low melt agarose in medium on a glass bottom dish with size 0 glass (IBL). Time-lapsed images were captured every 15 min from 5 hpi to 24 hpi.

### Image processing, cell counting and analysis

For cell counting images were processed using Fiji (14). Brightness and contrast were adjusted to identify eGFP, BrdU and Pax7 expressing cells. Cells were manually counted from z-stacks.

For processing of live imaging data, Z-projections were generated by maximum intensity projection and drift was corrected using the Fast4Dreg Plugin (15); the image was then manually cropped for cell segmentation. GFP positive cells were segmented using cellpose2 (16) with a custom trained model and cells tracked in Imaris 10.2 (Bitplane AG) using the Cells function with manual correction where necessary. Due to variations in the fluorescence intensity between samples, the threshold values were manually chosen for each individual sample to differentiate between visible cells and background noise. Following thresholding, the tracking algorithm was chosen to track the cells using following parameters: max distance = 5.00 µm, max gap size = 2 µm and track duration above 2.5 s.

Parameters of cell movement were exported for statistical analysis: distance from origin [µm], circularity, mean square displacement [µm^2^], speed mean [µm/s].

Neuromuscular junction (NMJ) structural integrity analysis was performed using NMJ Analyser as described by the authors (17).

### Statistical analysis

Statistical analysis was performed using GraphPad Prism Version 10.4.1. Measures of cell numbers acquired from samples processed by immunolabelling were tested for a normal distribution using a Kol-mogorov–Smirnov test. Significance was analysed using Student t-test. All p-values are indicated in the figures (ns: not significant; *p < 0.05; **p < 0.01; ***p < 0.001; ****p < 0.0001).

Multiple linear regression models with mixed effects were generated to test how mean square displacement (MSD) were affected by genotype or treatments in a time dependent manner using the Stata v15 package (18).

## Supplementary Figures

**Fig. S1.**
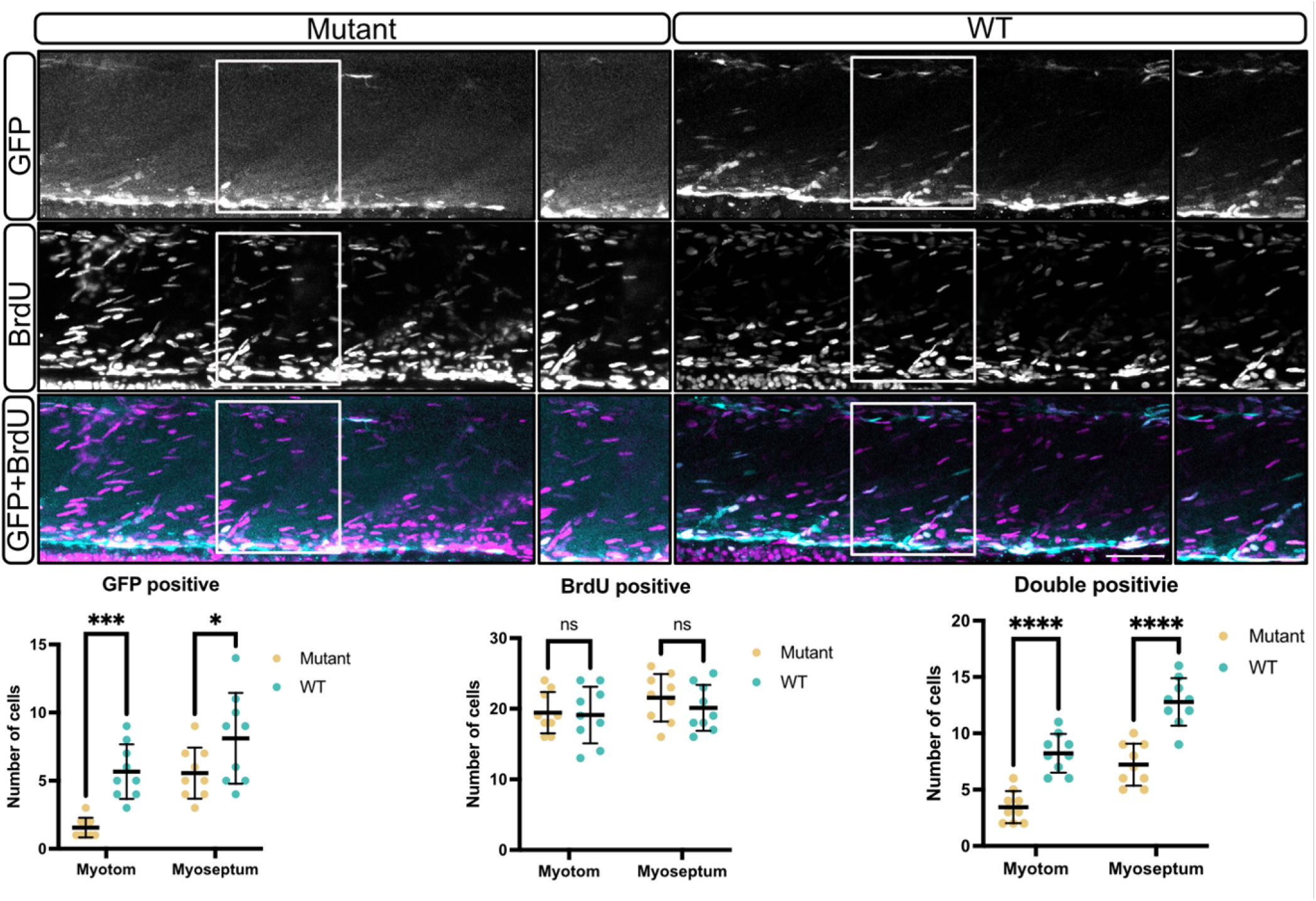
muSC proliferation is reduced in *tert* mutant zebrafish under homeostatic conditions. Representative images and quantification of muscle in 9 dpf uninjured *tert* mutant and WT larvae expressing Pax7a:egfp following 5 days BrdU exposure. Top panels show GFP (magenta, positive cells), middle panels show BrdU labelling (cyan; proliferating cells), and bottom panels display merged GFP/BrdU image. Quantification of GFP positive, BrdU positive, and GFP/BrdU double-positive cells was performed in both the myotome and myoseptum. Number of animals used n = 9 (all conditions). Data shown as mean ± SD and statistical testing performed using an unpaired Student’s t-test (ns not significant, * p < 0.05, *** p < 0.001, **** p < 0.0001) Scale bars: 50 µm.

**Fig. S2.**
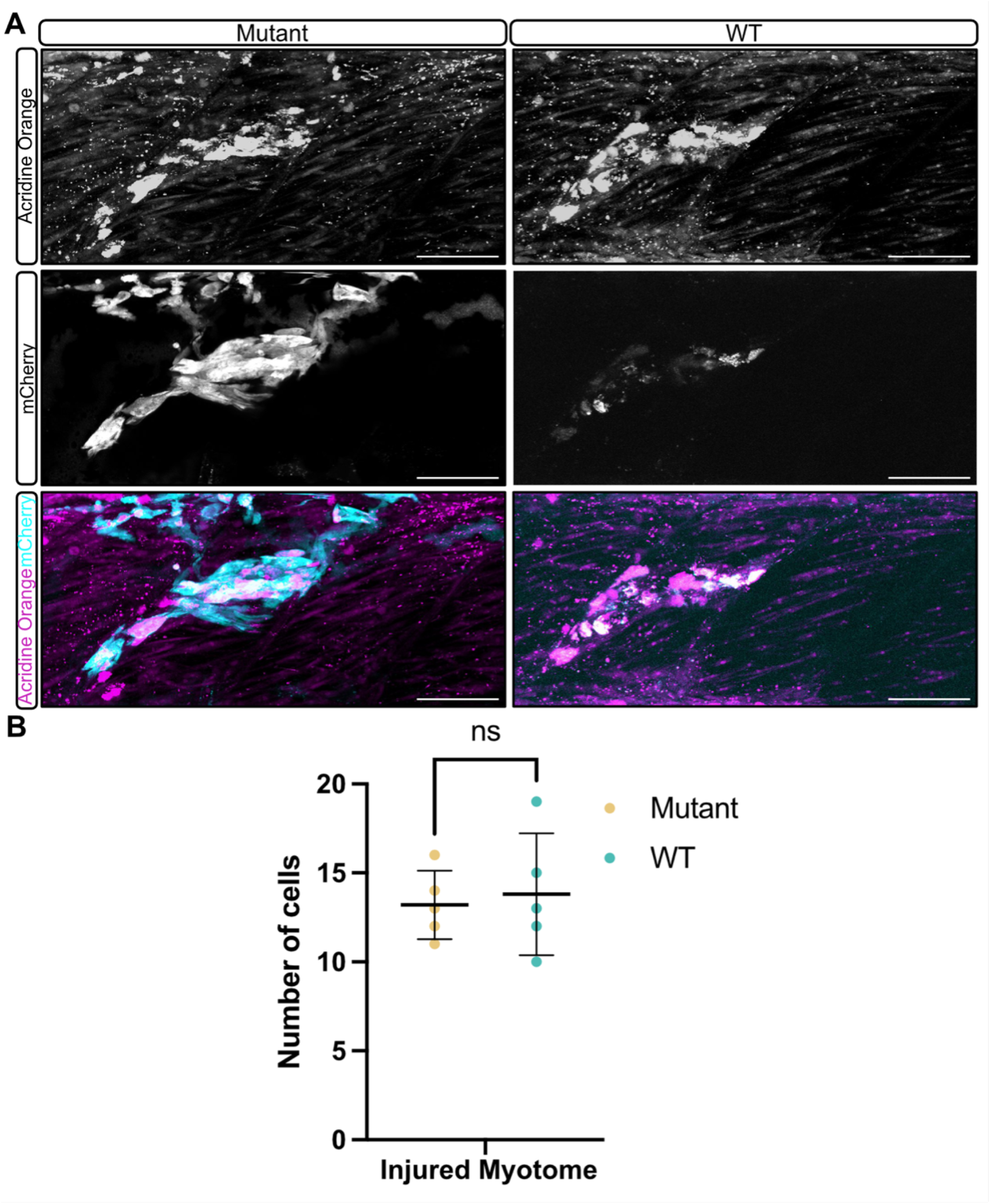
No significant difference in Acridine Orange positive cells in injured tert mutant and WT zebrafish muscle. (A) Representative images of injured myotomes in 5 dpf *tert* mutant and WT zebrafish larvae expressing fms:mCherry stained with Acridine Orange. (B) Quantification of Acridine Orange–positive cells in the injured myotome. Number of animals used n = 5 (all conditions). Data shown as mean ± SD and statistical testing performed using an unpaired Student’s t-test (ns not significant). Scale bars: 50 µm.

**Fig. S3.**
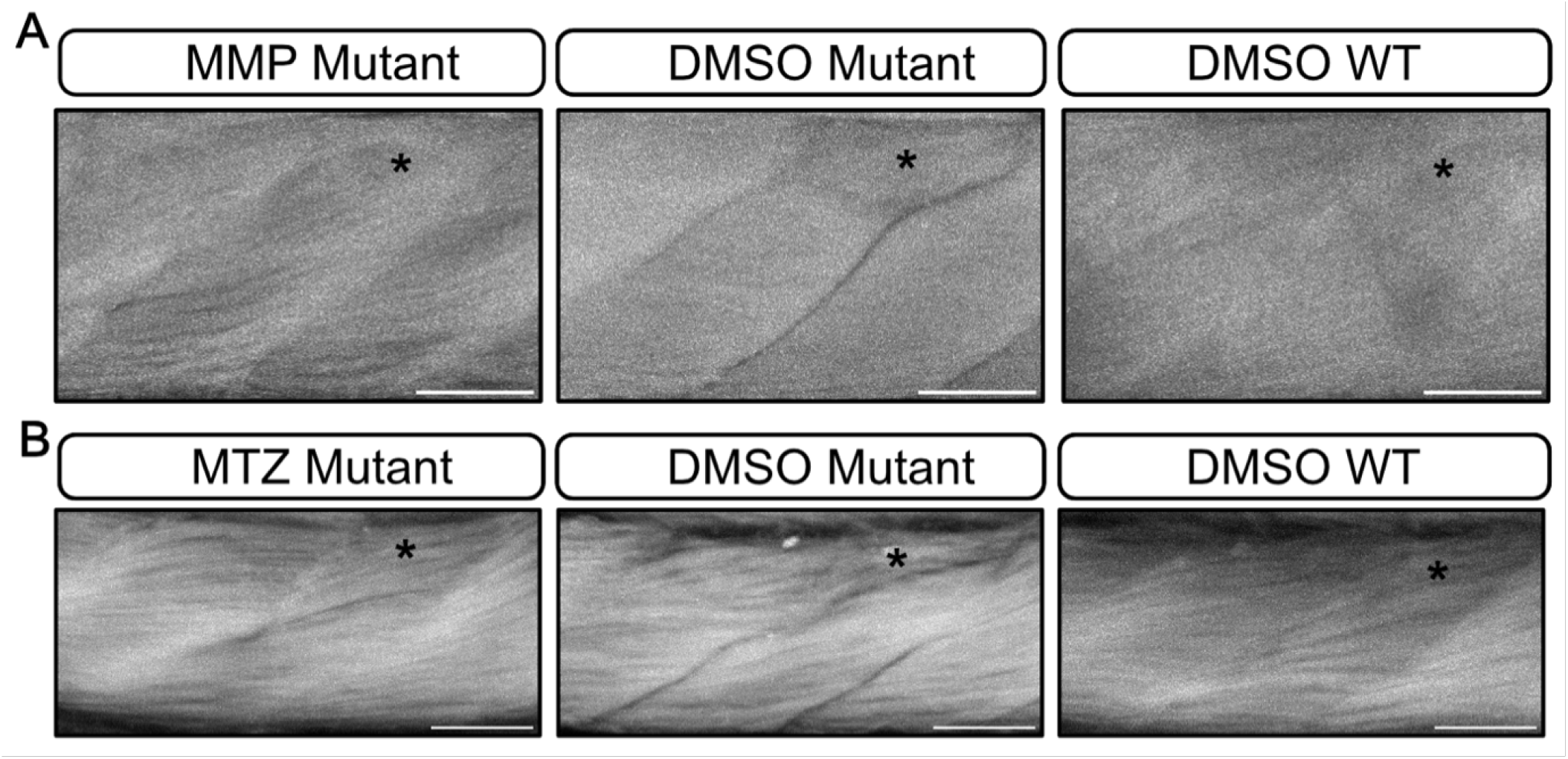
Muscle fibres alignment in MMP 9/13 inhibitor I treated and MTZ treated *tert* mutant following injury. (A–B) Representative phalloidin labelling of muscle at 6 dpi, showing myofiber morphology in larvae. (A) *tert* mutants treated with MMP9/13 Inhibitor I or DMSO, or WT treated with DMSO. (B) *tert* mutants treated with MTZ, *tert* mutant treated with DMSO and WT treated with DMSO. Asterisks mark the injury site. Scale bars: 50 µm.

**Fig. S4.**
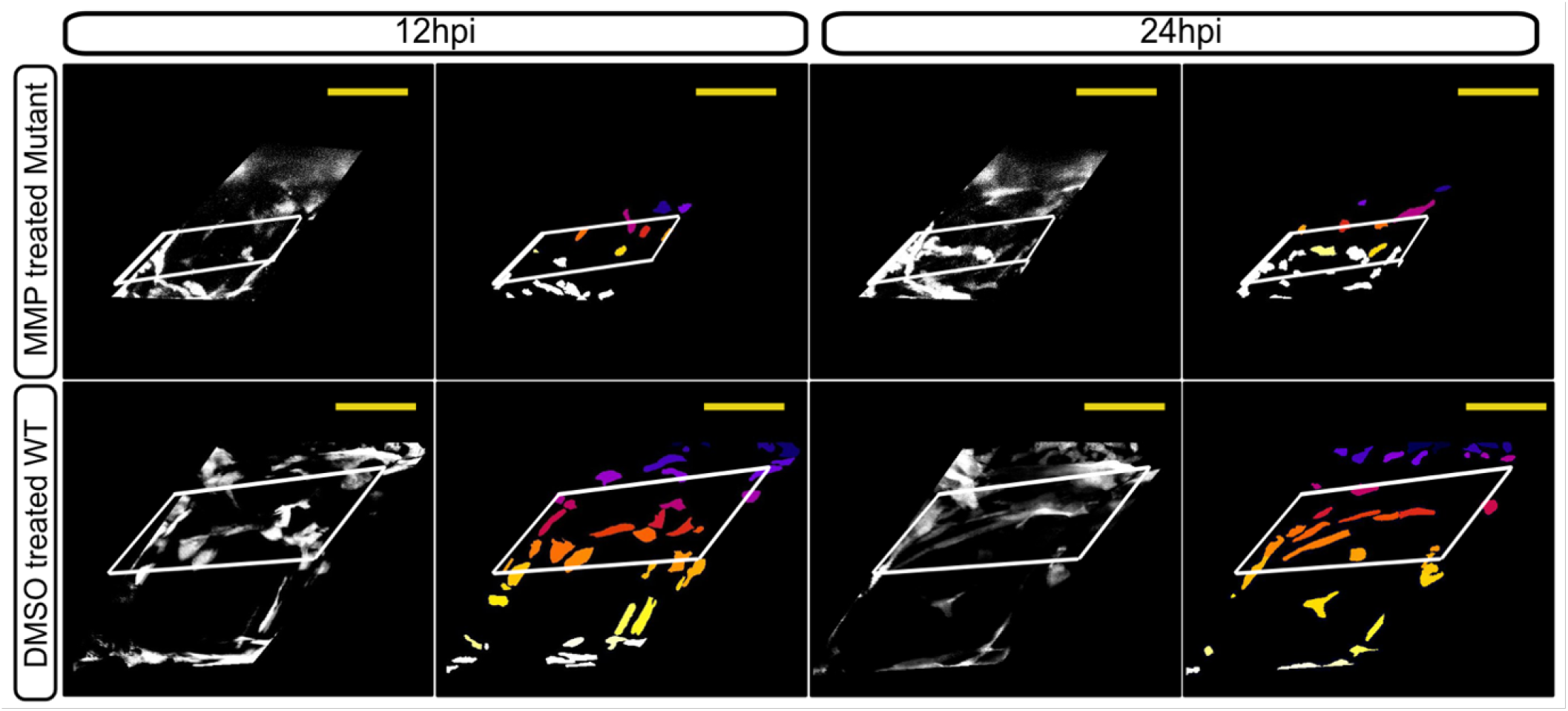
MMP9/13 inhibition enhances muSC migration to injured muscle of *tert* mutants. Representative images from time-lapsed recordings and corresponding segmented images of GFP positive muSCs in injured muscle of larvae expressing pax7a:egfp at 12 and 24 hpi. 5 dpf tert mutants were treated with MMP9/13 Inhibitor I 24 hours prior to injury and WT larvae were treated with DMSO. Scale bar 50 µm.

**Fig. S5.**
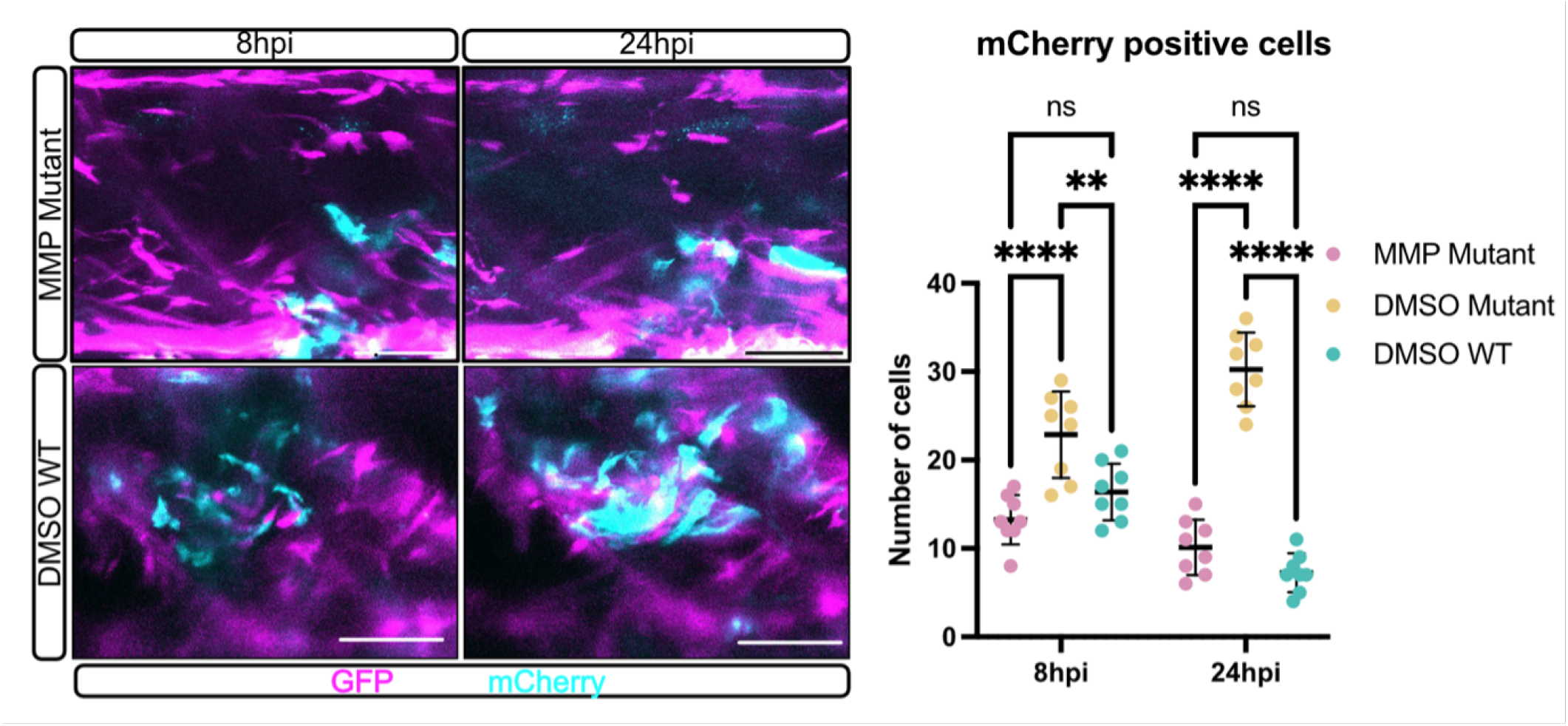
MMP9/13 inhibition reduces general macrophage accumulation at the injury site in *tert* mutant larvae. Representative live images (left) and quantification (right) of macrophages in injured muscle of *tert* mutant and WT zebrafish larvae treated with MMP9/13 inhibitor I or DMSO (control) at 8 hpi and 24 hpi. Images of muSCs expressing pax7a:egfp (magenta) and macrophages expressing fms:mCherry (cyan) were captured by time-lapsed imaging. Number of animals used n = 10 (*tert* mutants treated with MMP Inhibitor I), n= 8 (*tert* mutants treated with DMSO), n = 8 (WT treated with DMSO). Data shown as mean ± SD and statistical testing performed using an unpaired Student’s t-test (ns not significant, ** p < 0.01, *** p < 0.001, **** p < 0.0001). Scale bars: 50 µm.

**Fig. S6.**
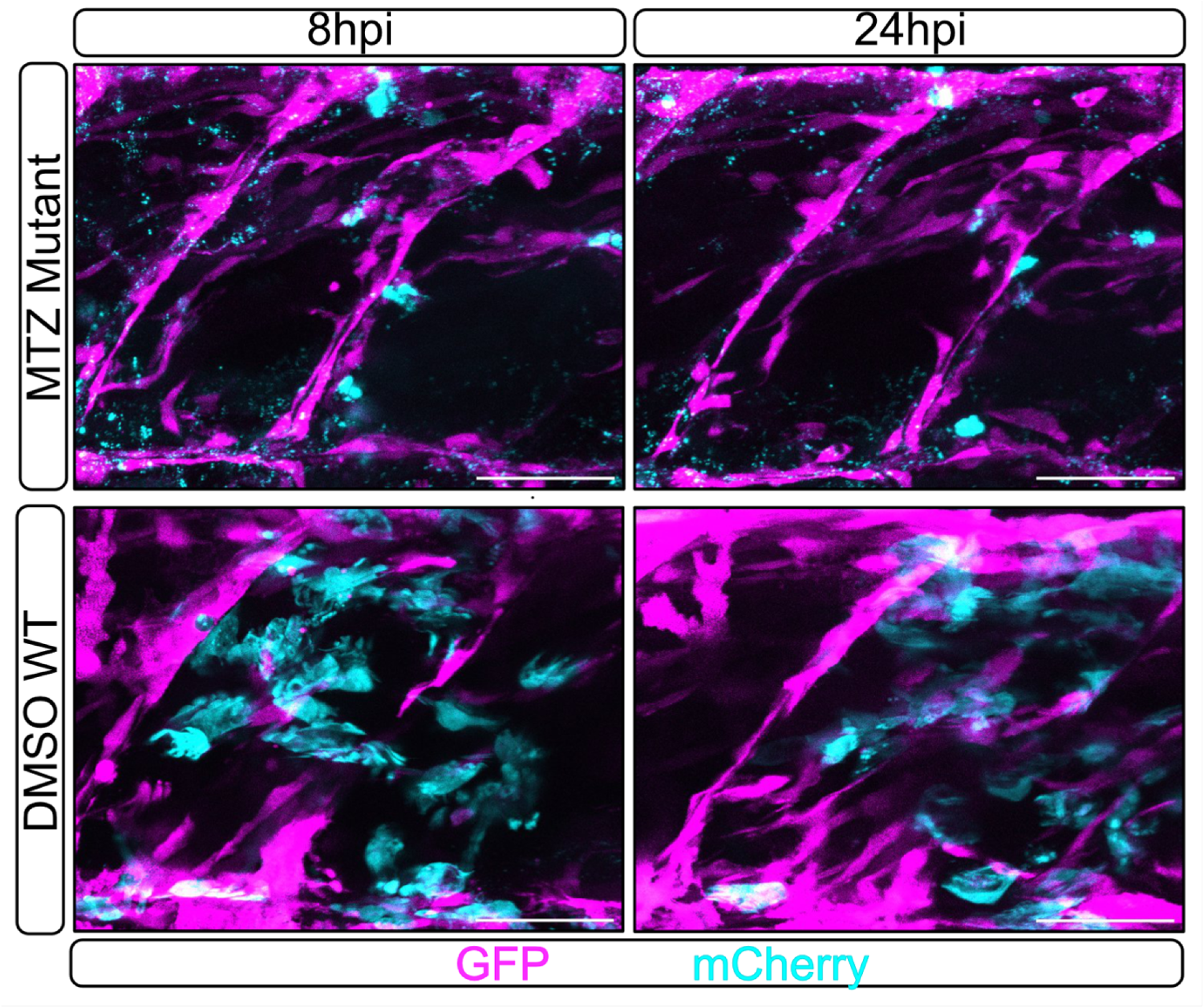
Evaluation of macrophage depletion in MTZ treated larvae. Representative images of *tert* mutant and WT larvae expressing pax7a:egfp (GFP, magenta) and fms:mCherry (mCherry, Cyan) transgenes at 8 and 24hpi after treatment with MTZ (*tert* mutant) or DMSO (WT). Scale bars: 50 µm.

**Fig. S7.**
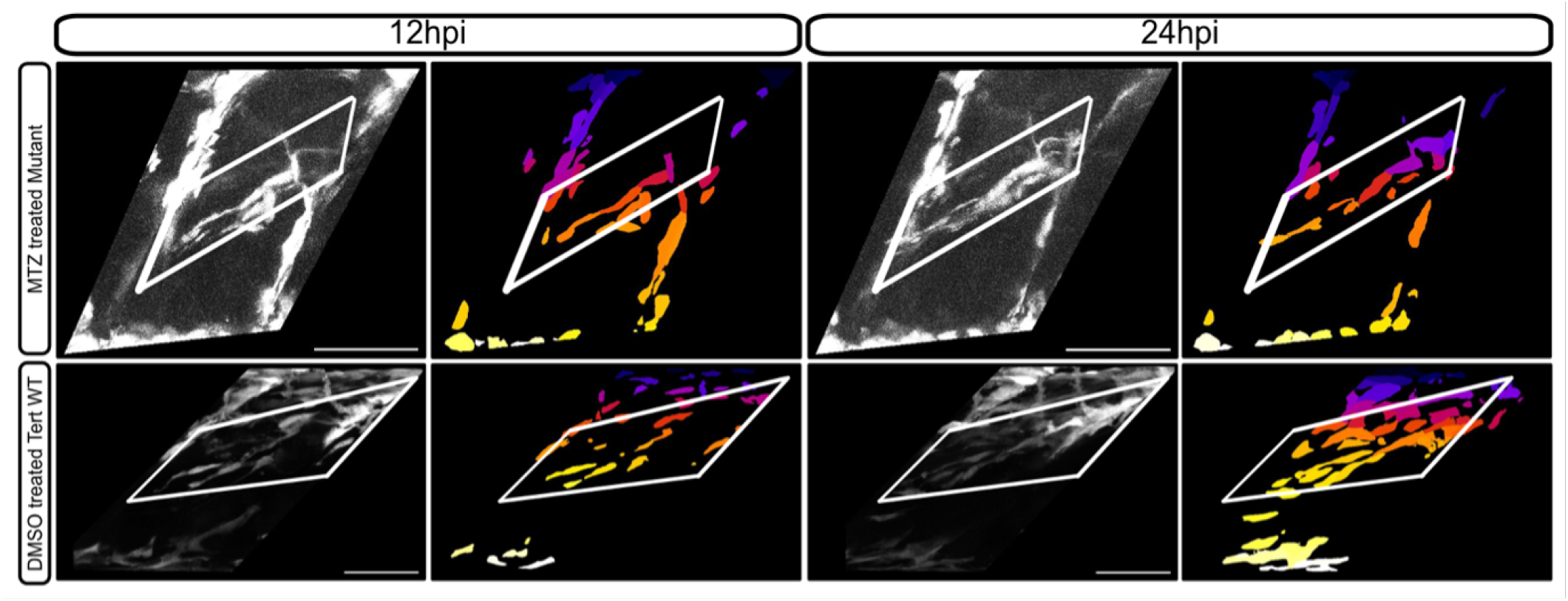
macrophage ablation rescues muSC migration to the injury site in *tert* mutants. Representative images from time-lapsed recordings and corresponding segmented images of GFP positive muSCs in injured muscle of larvae expressing pax7a:egfp at 12 and 24 hpi. 5 dpf *tert* mutants expressing fms:mCherry were treated with MTZ 24 hours prior to injury and WT larvae were treated with DMSO. Scale bar 50 µm.

